# A data-driven geospatial workflow to improve mapping species distributions and assessing extinction risk under the IUCN Red List

**DOI:** 10.1101/2020.04.27.064477

**Authors:** Ruben Dario Palacio, Pablo Jose Negret, Jorge Velásquez-Tibatá, Andrew P. Jacobson

## Abstract

Species distribution maps are essential for assessing extinction risk and guiding conservation efforts. Here, we developed a data-driven, reproducible geospatial workflow to map species distributions and evaluate their conservation status consistent with the guidelines and criteria of the IUCN Red List. Our workflow follows five automated steps to refine the distribution of a species starting from its Extent of Occurrence (EOO) to Area of Habitat (AOH) within the species range. The ranges are produced with an Inverse Distance Weighted (IDW) interpolation procedure, using presence and absence points derived from primary biodiversity data. As a case-study, we mapped the distribution of 2,273 bird species in the Americas, 55% of all terrestrial birds found in the region. We then compared our produced species ranges to the expert-drawn IUCN/BirdLife range maps and conducted a preliminary IUCN extinction risk assessment based on criterion B (Geographic Range). We found that our workflow generated ranges with fewer errors of omission, commission, and a better overall accuracy within each species EOO. The spatial overlap between both datasets was low (28%) and the expert-drawn range maps were consistently larger due to errors of commission. Their estimated Area of Habitat (AOH) was also larger for a subset of 741 forest-dependent birds. We found that incorporating geospatial data increased the number of threatened species by 52% in comparison to the 2019 IUCN Red List. Furthermore, 103 species could be placed in threatened categories (VU, EN, CR) pending further assessment. The implementation of our geospatial workflow provides a valuable alternative to increase the transparency and reliability of species risk assessments and improve mapping species distributions for conservation planning and decision-making.

## Introduction

An unprecedented rate of extinction is affecting millions of species worldwide (IPBES 2019). Hence, there is an urgent need to assess threats across species ranges to guide planning and priority-setting where conservation actions are most needed (Bachman et al. 2019). The International Union for Conservation of Nature (IUCN) Red List of Threatened Species employs two spatial metrics under criterion B (Geographic Range) to assess species extinction risk (IUCN 2012): Extent of Occurrence (EOO) and Area of Occupancy (AOO). These metrics represent the upper and lower bounds of a species distribution and have a different theoretical basis — the former measures the degree of risk spread and the latter is closely linked to population size (Gaston & Fuller 2009). However, the IUCN has recommended that for planning conservation actions, other metrics might be more appropriate (IUCN/SSC 2018; IUCN Standards and Petitions Committee 2019).

Recently, Brooks et al. (2019) suggested that IUCN Red List assessments should measure Area of Habitat (AOH), also known as Extent of Suitable Habitat. AOH is relevant to guide conservation by quantifying habitat loss and fragmentation within a species range, and is already part of the criteria for identifying Key Biodiversity Areas (KBA) (IUCN 2016a; KBA Standards and Appeals Committee 2019). To calculate AOH, researchers have relied mostly on refining expert-drawn range maps by clipping unsuitable areas based on published elevational limits and known habitat preferences (Harris & Pimm 2008; Ocampo-Peñuela et al. 2016). This approach has been used to inform the conservation status of different groups of terrestrial organisms on a global scale (Rondinini et al. 2011; Ficetola et al. 2015; Tracewski et al. 2016). However, expert-drawn range maps can have low accuracy due to errors of omission (known presences outside of the mapped area) and commission (known absences inside the mapped area) (Beresford et al. 2011; Peterson et al. 2018; Mainali et al. 2020). Generating range maps that minimize these errors is essential to avoid mischaracterization of species distributional patterns for local scale applications (Hurlbert & Jetz 2007) and to better inform conservation planning and decision-making (Rahbek 2005; Ficetola et al. 2014; Mainali et al. 2020)

There are different methods available to produce species ranges as an alternative to expert-drawn range maps, and it is crucial to understand the limitations and trade-offs when considering alternatives (Graham & Hijmans 2006; Cantú-Salazar & Gaston 2013; Maréchaux et al. 2017). One alternative is using species distribution models (SDM) that correlate environmental variables with known occurrences (Peterson et al. 2011). These methods have been successfully applied to estimate species ranges and inform Red List assessments (Pena et al. 2014; Syfert et al. 2014; Breiner et al. 2017). However, SDMs can be complex and involve methodological choices that if not well understood or explicitly communicated, can reduce transparency and reproducibility (Guisan et al. 2013; Sofaer et al. 2019; Feng et al. 2019). Furthermore, these models usually require species-specific fine-tuning of parameters (Radosavljevic & Anderson 2014; Muscarella et al. 2014) which makes them challenging to scale for hundreds (or more) species. Thus, there is an active search for more straightforward approaches (García□Roselló et al. 2019).

Spatial interpolation methods are an alternative approach to support the mapping of species distributions and have been widely employed for conservation applications (Li & Heap 2008). Of the different types of interpolation procedures available, Inverse Distance Weighting (IDW) is a deterministic method that is accessible, user-friendly, and could be readily employed to support producing species ranges. IDW uses known presences and absences to produce a surface of probability values for the occurrence of a species within a given area (Hijmans & Elith 2019). The main assumption is that species are more likely to be found closer to presence points and farther from absences (*i.e.* spatial autocorrelation), and the local influence of points (weighted average) diminishes with distance. IDW has been used for mapping invasive plants (Roberts et al. 2004) and to understand the distribution of coral reef sessile organisms, with a higher accuracy than more advanced geostatistical interpolation methods (Zarco-Perello & Simões 2017). Recently, Gomes et al. (2018) found that Maxent SDMs models largely overlapped with abundance maps derived from using IDW based inventory plots for 227 hyper-dominant Amazonian tree species. Furthermore, IDW may outperform SDM’s when environmental variables do not provide more explanatory power than the spatial structure of the occurrence data (Bahn & McGill 2007; Hijmans 2012). Thus, IDW is also useful to compare it to other approaches in species distribution modeling (Raes & ter Steege 2007; Hijmans 2012; Hijmans & Elith 2019).

Here, we developed a geospatial workflow to map species distributions with presence and absence points derived from primary biodiversity data. We aimed to provide a reproducible and transparent approach to (1) improve the accuracy of species ranges and the estimation of Area of Habitat (AOH) for local-scale applications and (2) increase the reliability of extinction risk assessments under the IUCN Red List. To show the advantages of our approach we implemented our workflow on 2,273 bird species in the Americas, 55% of all terrestrial birds found in the region. Birds represent a standard in the evaluation of species conservation status (BirdLife International 2018) and comprise more than half of the occurrence records on biodiversity databases (Troudet et al. 2017). Overall, by highlighting differences in produced ranges and derived spatial metrics for a well-known group of organisms in a species-rich region, we present the case for an improved approach at mapping species distributions for conservation planning and decision-making.

## Methods

### Geospatial workflow

We developed a data-driven, reproducible geospatial workflow (Fig. 1) to refine the distribution of a species starting from its Extent of Occurrence (EOO) to producing a final Area of Habitat (AOH) map. First, the user must gather and process occurrence data, an essential pre-processing step of any range mapping procedure. We suggest users follow recent guidelines e.g. (Araújo et al. 2019; Feng et al. 2019) to that end. Our proposed method consisted of five main steps:

**Figure 1.**
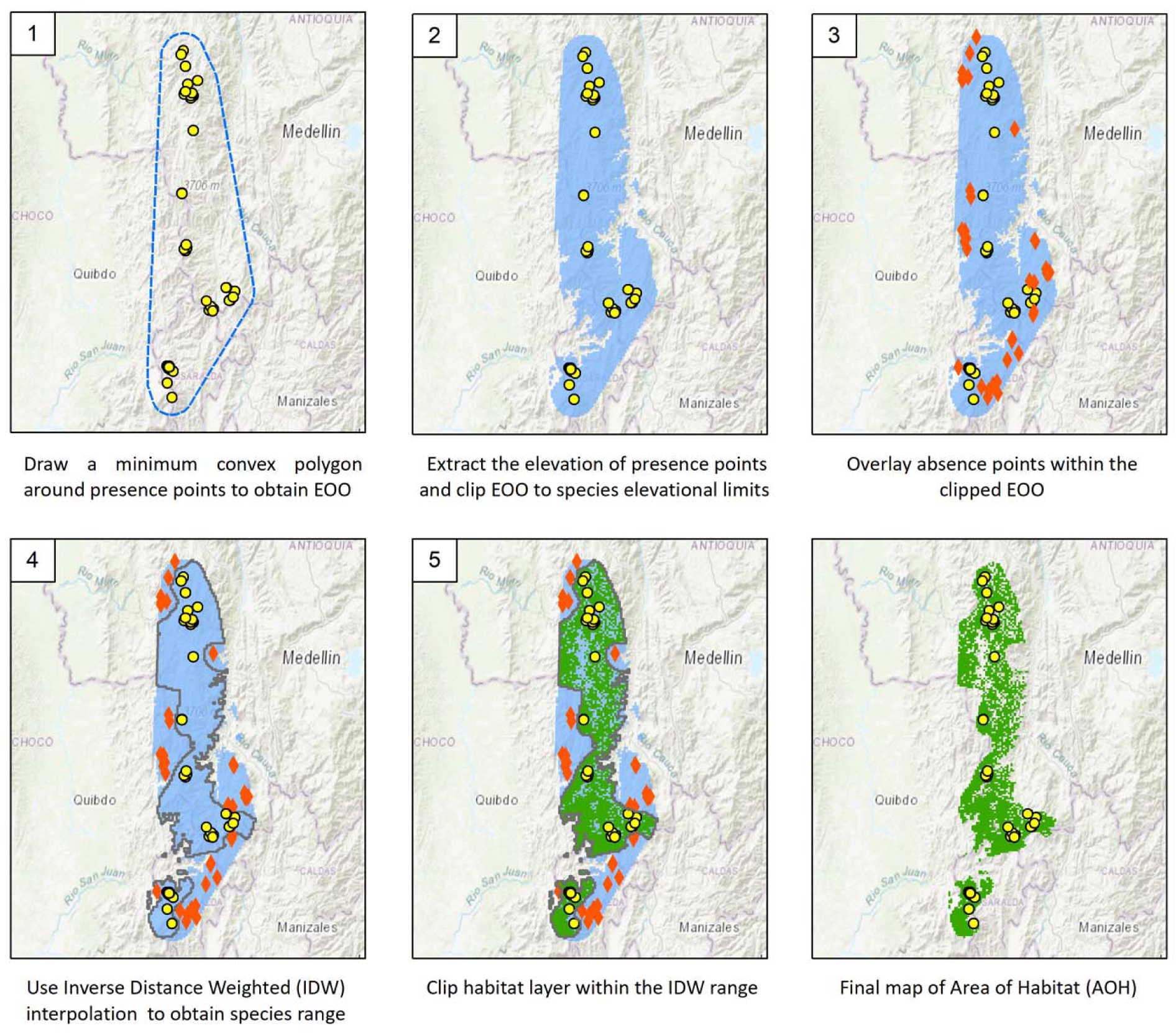
Mapping protocol to refine the distribution of a species from Extent of Occurrence (EOO) to Area of Habitat (AOH). This is an illustrative example for the Glittering Starfrontlet (*Coeligena orina*), a hummingbird species in the cloud forests of the western Andes of Colombia (Fig. 2 for species picture). The EOO was buffered by 10 km following the implementation in our workflow (see methods).

1. Draw the Extent of Occurrence (EOO) around presence points. The EOO is the upper bound of the distribution of a species (Brooks et al. 2019) and should be mapped as the Minimum Convex Polygon (MCP) due to its consistency and scale-free properties (Joppa et al. 2016; IUCN Red List Technical Working Group 2019; IUCN Standards and Petitions Committee 2019).
2. Clip the EOO to the elevational limits of a species by extracting the elevation of each occurrence point using a Digital Elevation Model (DEM). The choice of DEM resolution depends on the spatial extent of the data and the intended analysis for which 90 m to 250 m resolutions are commonly employed (Amatulli et al. 2018)
3. Overlay absence points to the clipped EOO. Absences can be derived from known surveys, monitoring schemes, or citizen science projects that have a measure of sampling effort such as eBird or biodiversity atlases (Johnston et al. 2019).
4. Interpolate presences and absence points using Inverse Distance Weighting (IDW). The output is a raster surface within the clipped EOO where each cell has a value between 0 and 1. The species range is obtained by using a threshold to convert continuous results to a binary score where the species is either present or absent (Liu et al. 2005). The ranges produced with IDW are consistent with the observation that distributions are naturally porous and have discontinuities (holes) at higher resolutions (Hurlbert & Jetz 2007).
5. Clip a habitat layer inside the species IDW range to remove unsuitable areas and derive Area of Habitat (AOH) (KBA Standards and Appeals Committee 2019). We recommend following the IUCN Habitat Classification Scheme for consistency. Users could use empirically developed products (e.g., Nature Map Explorer – www.explorer.naturemap.earth) or use expert criteria to select appropriate landcover classes that match the definition of species habitats coded by the IUCN.

### Case Study

Here we show the implementation of our proposed geospatial workflow using data for terrestrial bird species in the Americas. We followed the Handbook of the Birds of the World-Birdlife Taxonomic Checklist v4, which is the basis for the IUCN Red List of Threatened Species and bird species distribution maps of the world (HBW and BirdLife International 2019). We selected bird species with ranges smaller than the global median [< 1M km^2^ (Orme et al. 2005; Jenkins et al. 2013)] because they are typically more at risk of extinction (Chichorro et al. 2018). We excluded species endemic to the Galapagos and Hawaiian Islands and those classified as Extinct (EX), Extinct in the Wild (EW), and possibly extinct (CR-PE) according to the 2019 IUCN Red List. Our final dataset comprised 2,273 bird species in the Americas, 55% of all terrestrial birds found in the region (BirdLife International 2020). We generated ranges for all species and derived Area of Habitat (AOH) for a subset of 741 species whose only habitat type is “forest” according to the IUCN Habitat Classification Scheme (Donald et al. 2018).

#### 1. Primary biodiversity data

##### 1.1 Gathering and processing occurrence data

Occurrence data was gathered from the Global Biodiversity Information Facility (GBIF; https://www.gbif.org). The GBIF biodiversity database is a repository of multiple occurrence datasets spanning from museum specimen records to the eBird and iNaturalist citizen science projects (GBIF 2020). We obtained occurrence data with the R package *rgbif* (Chamberlain & Boettiger 2017) using the ‘occ_search’ function in April 2019. We removed duplicated coordinates and removed species with five records or less (Pender et al. 2019). We used the R package *CoordinateCleaner* (Zizka et al. 2019) to automatically filter common erroneous coordinates in public databases such as those assigned to the sea, country capitals, or biodiversity institutions (Maldonado et al. 2015).

The GBIF occurrence data operates on a stable taxonomic backbone that although useful for database management does not represent current knowledge on species limits (Burfield et al. 2017). Thus, we consulted the Avibase database (https://avibase.bsc-eoc.org; Lepage et al. 2014) to check for species that differed from the HBW-Birdlife taxonomy, and manually updated their occurrence data with the interactive modular R platform *Wallace* (Kass et al. 2018). In this process we accounted for taxonomic splits and removed obvious geographical outliers that we deemed as occurrence points in areas where a species is not able to disperse given the presence of large geographical barriers (Barve et al. 2011; Hazzi et al. 2018).

##### 1.2 Deriving absences

We detected absences based on eBird hotspots – publicly accessible locations that are frequently birded and have been previously approved by an administrator. Here, we deemed absences as eBird hotspots where a given species *has never been recorded.* We used this approach as a stringent criterion to minimize the uncertainties regarding absences (Lobo et al. 2010). We used the R package *auk* (Strimas-Mackey et al. 2016) to extract from the eBird Basic Dataset (2019) observational checklists for 57 countries and territories in the Americas. We limited the data to complete checklists (birding events where observers report all species detected) from January 1, 2000 to April 31, 2019. Using complete checklists is a critical step to obtain better models of species distribution and is recommended as part of the eBird best practices (Johnston et al. 2019; Strimas-Mackey et al. 2020). We obtained 142,703 points locations corresponding to eBird hotspots which we used to identify absences (see mapping protocol).

#### 2. Detailed Workflow

The implementation of our workflow followed methodological choices that were conceived to work for a large number of species and computational efficiency, mostly using default parameters that users can adapt or modify depending on the intended purpose. Our workflow was run on CyVerse’s Atmosphere cloud-computing platform to speed processing time (see acknowledgments). Following best practices for sharing large datasets (Marwick et al. 2018), we provide the R code to reproduce it on a small sample of species (see Code Availability statement).

In our procedure we drew each species EOO and then buffered it to standard 10 km (Ocampo-Peñuela & Pimm 2014). Then, we extracted elevational limits from occurrence points using the SRTM v4.1 DEM at 250 m resolution (Jarvis et al. 2008) which is a good compromise between computational ease and resolution in complex topographical areas such as mountains (Hazzi et al. 2018). The interquartile range (IQR) was used as a robust test for outlier detection, in which elevational data points below Q_25_ − 1.5 *IQR, or above Q_75_ + 1.5 *IQR were removed (Wilcox 2017). We overlaid absence points to the clipped EOO and removed those within a 5 km buffer of any presence points (Johnston et al. 2019; Strimas-Mackey et al. 2020).

We computed IDW in the R package *dismo* (Hijmans et al. 2017) using all presence and absence points available for each species. To derive AOH for forest species we used a recently developed 50-m global forest/non-forest map from the TanDEM-X satellite (Martone et al. 2018). This product is designed to specifically estimate forest cover instead of other approaches that are more focused on evaluating forest cover change [*e.g.* Hansen et al. (2013)]. We aggregated to 250 m with a nearest neighbor assignment to match the resolution of our elevational data.

#### 3. Comparison of species ranges

We compared the IDW and BirdLife ranges in terms of accuracy, range size, and range overlap. We also evaluated differences in the estimation of Area of Habitat (AOH). For the BirdLife ranges, we examined how accuracy influences their range size and the amount of spatial overlap with the IDW ranges.

##### 3.1 BirdLife distribution maps

We obtained the BirdLife distribution maps v.2019-1 (created November 2019) that come assembled in an ESRI File Geodatabase (http://datazone.birdlife.org/species/requestdis). Each species is represented by polygons coded by different categories of presence, origin, and seasonality (BirdLife International and HBW 2019). We combined all coded polygons to represent each species range into a single spatial feature (Cantú-Salazar & Gaston 2013) using ArcGIS Pro 2.4.1. To estimate Area of Habitat (AOH) from the BirdLife ranges, we refined the polygon of each species by elevation and forest cover (*sensu* Ocampo-Peñuela et al. [2016]). We used the 50-m global forest/non-forest map that we employed in our geospatial workflow. Species elevational limits were obtained from the IUCN Red List and HBW Alive.

##### 3.2 Accuracy assessment

We evaluated the discriminant capacity of the IDW and BirdLife ranges to classify presences and absence points within each species EOO. We used three basic metrics of classification performance: errors of omission, errors of commission, and overall accuracy (Anderson et al. 2003). Omission errors were quantified based on the amount of GBIF presences left outside of the mapped range (false negatives), whereas commission errors were quantified based on the amount of eBird absences that remain inside the mapped range (false positives). The overall accuracy was calculated using Jaccard’s similarity index, which summarizes the ability of the mapped range to (a) maximize true presences, (b) reduce both false negatives and false positives, and (c) disregard true negatives that are easily classified due to issues of prevalence (Li & Guo 2013; Leroy et al. 2018).

The accuracy metrics were calculated with a confusion matrix that cross-tabulates the match between predictions and observations (Fig. 2 for reference). For the BirdLife ranges we computed the accuracy metrics overlaying all the available presences and absences of each species. For the IDW ranges, we conducted a *k*-fold cross-validation procedure using 80% of the data for training and 20% for testing the model, resulting in five different folds for evaluation (Gareth et al. 2013). The resulting confusion matrices of each fold were summed to represent the total performance of the model and compute the final accuracy metrics.

**Figure 2.**
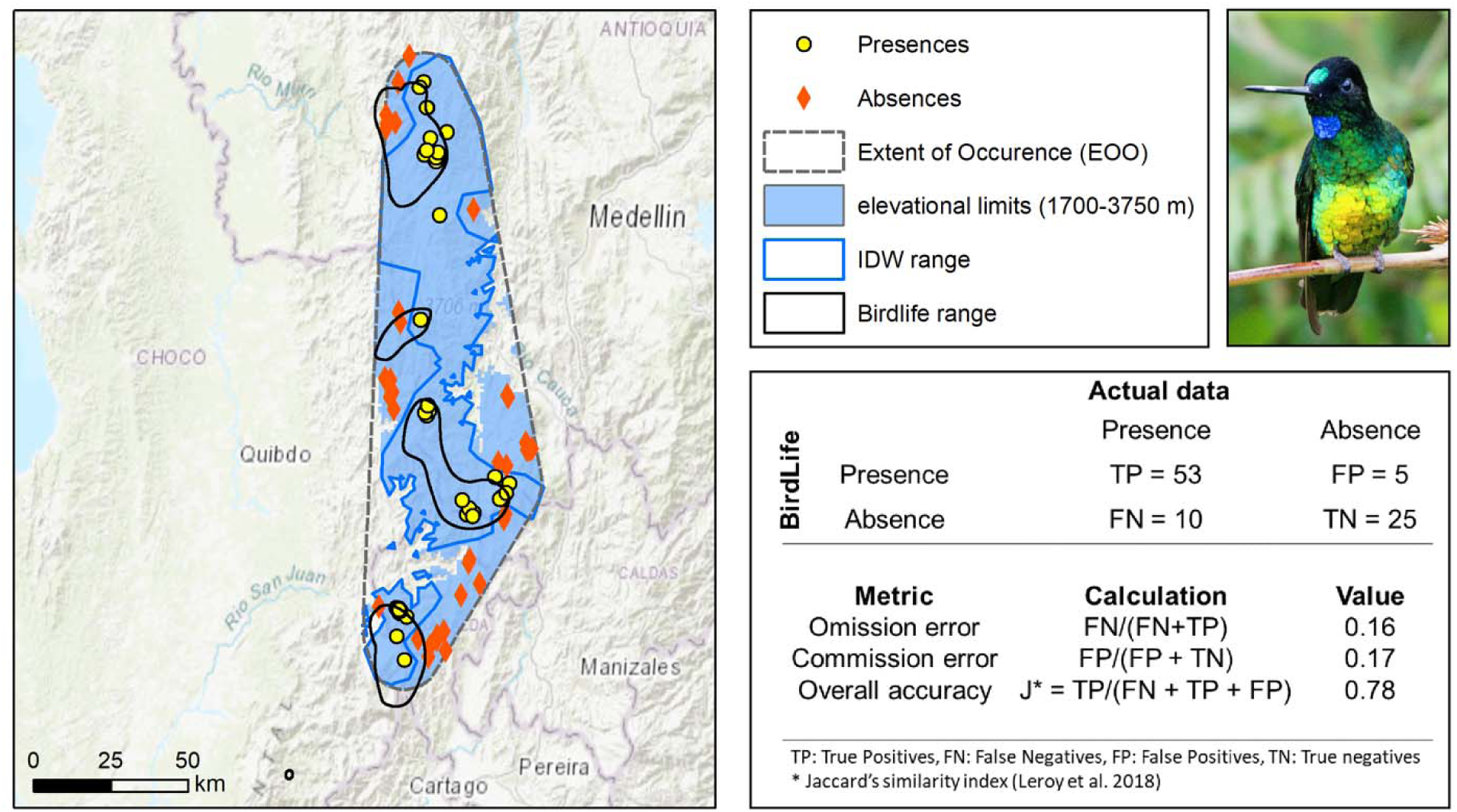
Example of accuracy analysis for the BirdLife range of the Glittering Starfrontlet (*Coeligena orina*) in the western Andes of Colombia, using a confusion matrix to calculate accuracy metrics. The overall accuracy was high (0.78) but note the arbitrary division of the range into four areas, including a fifth one indicated with the arrow in which no data points are available on GBIF (up to April 2020). This area did not enter into the accuracy calculation because it is outside of the EOO (see methods). The IDW range is shown for visual comparison. TP = True Presences; FP = False Presences; TN= True Negatives; FN = False Negatives. Photo by Anderson Muñoz.

##### 3.3 Range size comparison

We evaluated differences in range size between the IDW and BirdLife ranges for our complete dataset of 2,273 species. For this, the IDW prediction surface was transformed into a binary map (presence/absence). We used a probability threshold of > 0.5 to retain areas where the species is more likely to occur. This threshold is particularly useful to compare the areas of different distribution maps that have been produced for any given species (Zhang et al. 2005). We tested for a statistically significant difference between both datasets with a Two-sample Kolmogorov–Smirnov test (K-S test). Then, we quantified the differences in the distribution of range sizes across deciles with a shift function (Fig. 3) using the *rogme* package in R (Rousselet et al. 2017).

**Figure 3.**
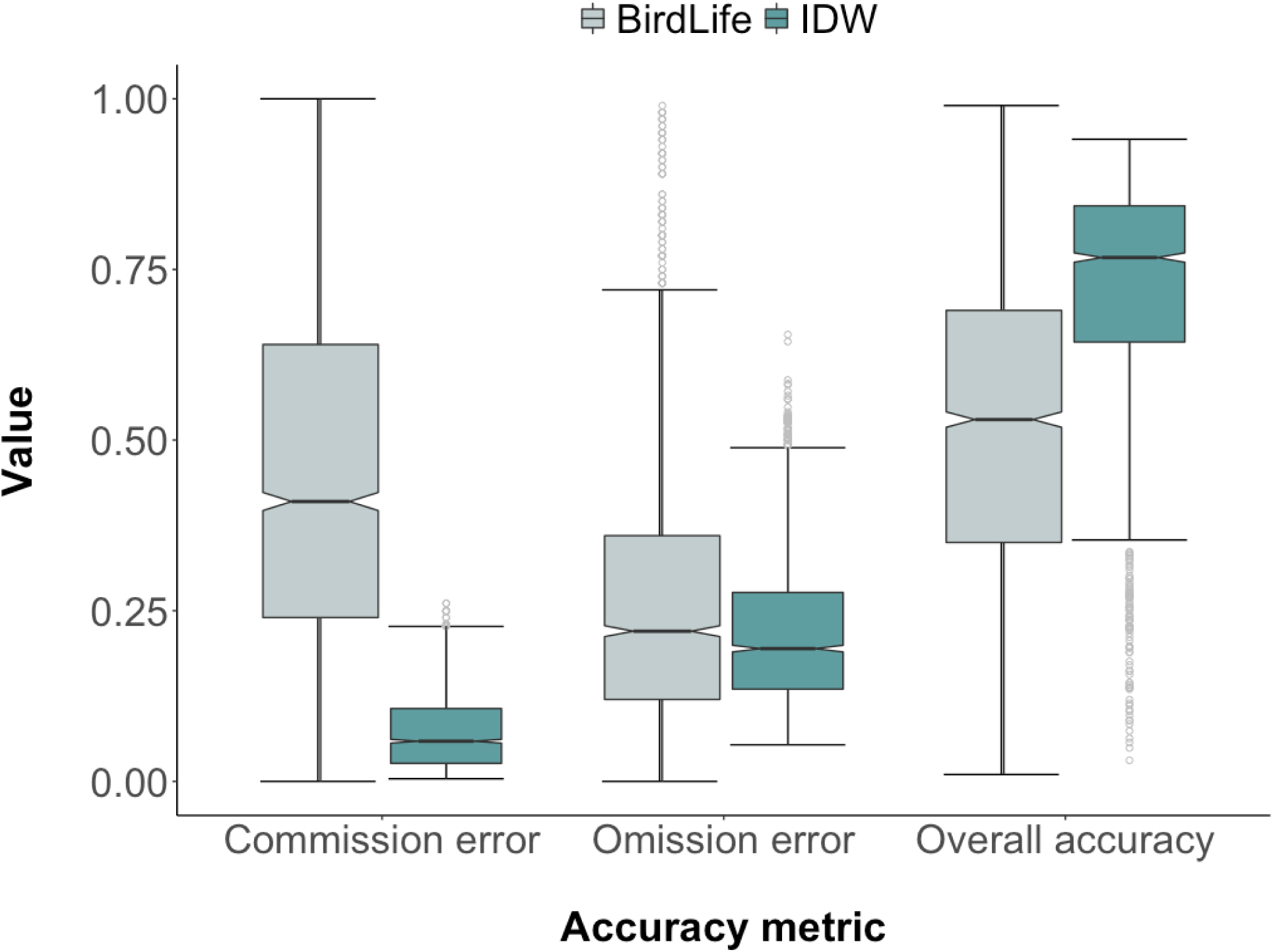
Values for each of the three evaluated metrics of accuracy (commission error, omission error and overall accuracy) between the IDW and the BirdLife ranges. The boxes show the Interquartile Range (IQR) which visualize the spread of 50% of the data points between the 25^th^ and 75^th^ percentile. The notches represent 95% confidence intervals of the median indicating differences because of no overlap. Note that most distributions are skewed, and data points outside of the whiskers (1.5 x IQR) are not necessarily outliers. There was a statistically significant difference between the values of the IDW and BirdLife ranges for each evaluated metric (two-sample K-S test, p < 0.001).

We also computed a robust log-linear regression with an MM-type estimator in the R package *robustbase* (Maechler et al. 2019) to examine how omission and commission errors influence difference in range size between IDW and BirdLife datasets. We further calculated the 20% trimmed mean of range size for each dataset to assess how the typical IDW and BirdLife ranges compare to each other. Trimmed means are a robust measure of centrality recommend to estimate average values of skewed distributions (Wilcox 2017). We estimated a 95% Confidence Interval (CI) of the range size difference using a percentile bootstrap procedure. The same approach was used to estimate differences in Area of Habitat (AOH) for our subset of 741 forest species. We also measured the effect size with a Brunner–Munzel test, which estimated the probability that a randomly sampled range from one of the datasets is larger than another randomly sampled range from the other dataset. Statistical analysis was conducted with R functions provided by R.R. Wilcox (https://dornsife.usc.edu/labs/rwilcox/software)

#### 4. Comparison of IUCN extinction risk assessments

We used the R package *ConR* (Dauby et al. 2017) to compute with the gathered occurrence data, apreliminary extinction risk assessment based on the IUCN criterion B [Geographic Range] parameters(IUCN 2012): Extent of Occurrence (EOO) and Area of Occupancy (AOO). Species were assigned to threatened categories when threshold levels are met for either EOO or AOO (e.g. AOO: Vulnerable<2,000 km^2^). Note that AOO is calculated by tallying the area of 2 x 2 km grid cells with documented presences (IUCN Standards and Petitions Committee 2019). Thus, it is a subset of Area of Habitat (AOH) (Brooks et al. 2019). In addition to EOO or AOO thresholds, at least two out of three other conditions must be met if a species is to be assigned to threatened categories (IUCN 2012). Two of these conditions are continued decline in habitat and number of locations, defined as areas in which a single threatening event can affect all individuals of a species (IUCN Standards and Petitions Committee 2019).

*ConR* assumes that habitat is declining for all species and quantifies locations based on 10 km^2^ gridded squares around presence data, for which 10 or fewer locations are the threshold to meet threatened categories (IUCN 2012). These assumptions to automate the process help assessors to find and prioritize species and should be interpreted as a first filter for a full IUCN extinction risk assessment.

We used the R package *rredlist* (Chamberlain 2019) to query from the IUCN Red List 2019 then following data: species threat categories and criteria, upper and lower elevational limits, EOO and AOO metrics. We compared differences in EOO and AOO measurements between our automated protocol and the Red List. Then we evaluated the agreement between the threatened categories (Vulnerable -VU, Endangered -EN, Critically Endangered - CR) in the preliminary automated assessment and the IUCN Red List 2019 using Kendall’s coefficient of concordance (Kendall’s *W*) which ranges between 0 and 1 (no agreement to complete agreement).

## Results

### 1. Accuracy

We found that for the complete dataset of 2,273 bird species, our ranges had reduced errors of omission and commission with increased overall accuracy within the species EOO (Fig. 3). For the BirdLife ranges, the 20% trimmed mean of omission errors was 0.23 (95% percentile bootstrap CI [0.22; 0.23], p < 0.001). Commission errors were common (0.43; CI: 0.41-0.44, *p* < 0.001) and the overall accuracy of the ranges was only moderate (0.52; CI: 0.51-0.53, *p* < 0.001). In comparison, For the IDW ranges omission errors were lower (0.20; CI: 0.19-0.21, p < 0.001). Commission errors were very low (0.06; CI: 0.06-0.07, p < 0.001) and the overall accuracy was higher (0.76; CI: 0.75-0.76, p < 0.001). All of these differences were statistically significant (Two-sided KS-test; *p* < 0.001). In percentages, 90% of our ranges had a higher overall accuracy than the BirdLife ranges, 52% had lower omission errors and 85% had lower commission errors.

### 2. Range size and spatial overlap

The IDW and BirdLife datasets had different distributions of range sizes for the 2,273 bird species analyzed (Fig 4A; K-S test = 0.18, *p* < 0.001). A shift function showed the BirdLife ranges are consistently larger than the IDW ranges across the entire distribution of values with positive differences between all deciles (Fig 4B).

**Figure 4.**
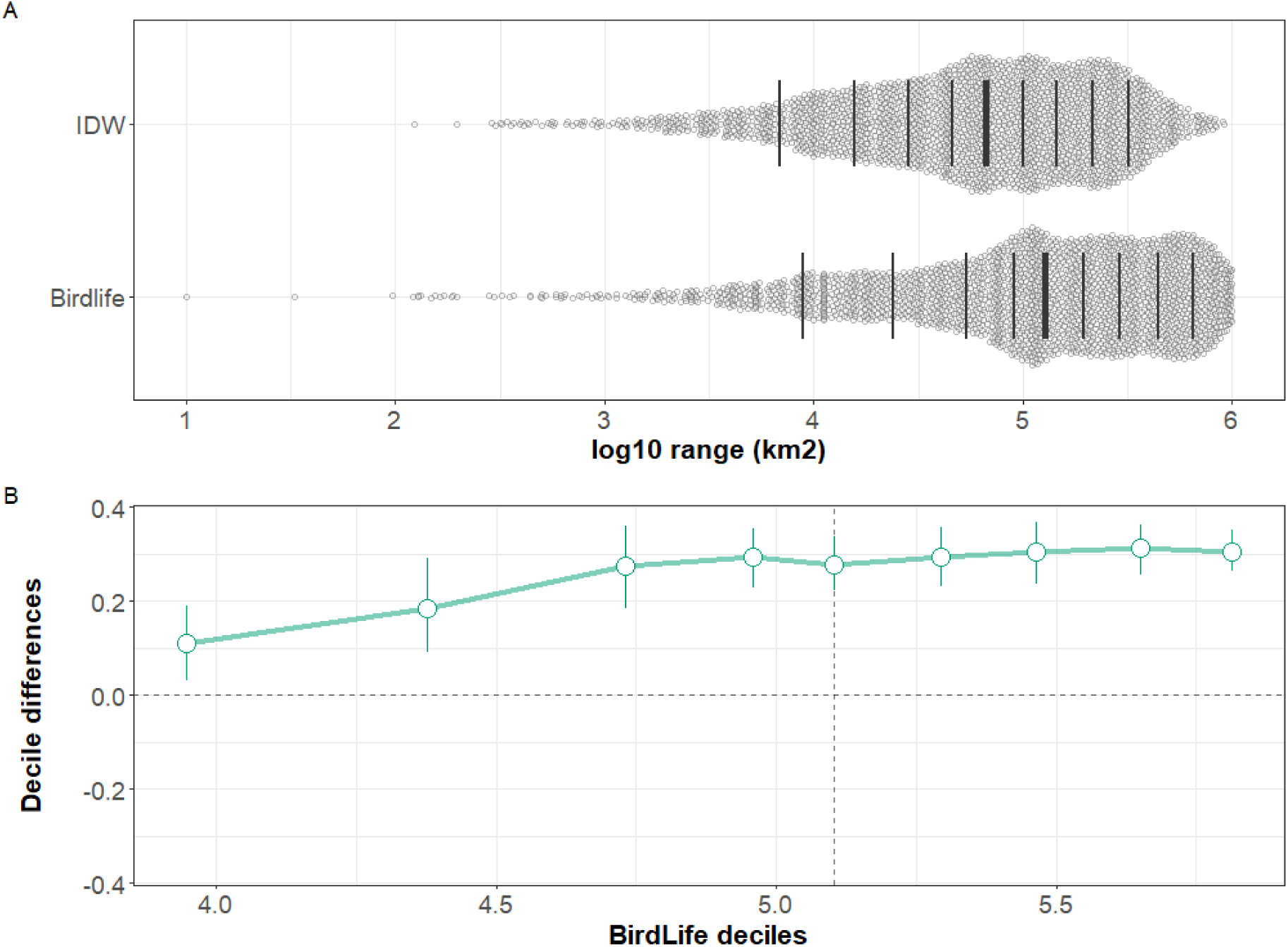
**A.** Comparison of the distribution of range sizes between IDW and BirdLife for 2,273 birds in the Americas. The thicker line corresponds to the median of the distribution and the thinner lines represent deciles **B.** Pairwise differences showed that for each decile across the distribution the BirdLIfe ranges are consistently larger in value than the IDW ranges. These difference are estimated with 95% confidence intervals derived using a percentile bootstrap procedure (Rousselet et al. 2017)

The 20% trimmed mean showed that the typical BirdLife range is almost twice as large as the typical IDW range (BirdLife _*tmean*_ = 163,809 km^2^; IDW _*tmean*_ = 82,843 km^2^) with an estimated difference of 80,966 km^2^ (CI:69,396 - 93,254 km^2^], *p* < 0.001). The percentage of BirdLife ranges that are larger than ours is 79% (n = 2,273). As a measure of effect size, the probability that a BirdLife range is larger than an IDW range for a given species if selected at random is 0.61 (CI 0.60 - 0.63, *p* < 0.001). The estimated Area of Habitat (AOH) derived from BirdLife ranges was also larger than those derived from the IDW ranges (BirdLife-AOH _*tmean*_ = 47,499 km^2^; IDW-AOH_*tmean*_ = 38,745 km^2^) with an estimated difference of 8,753 km^2^ (CI: 1,913 - 15,488 km^2^, *p* = 0.013).

We found that commission errors increased differences in range size for the BirdLife ranges. A robust log-linear regression showed that commission errors have a larger positive effect (*coef* = 1.24, *SE* = 1.14, *t* = 6.45, *p* <0.001) than the negative effect of omission errors (*coef* = −0.79, *SE* = 0.25, *t*= −3.13, *p* = 0.02) on range size difference (*intercept* = 10.40, 0.14, t = 74.47, p < 0.001). Both independent variables were significant predictors of difference in range size (*X*^*2*^ =9.82, *df* =1, *p* = 0.002). The spatial overlap between the BirdLife and IDW ranges was only 28% (95% percentile bootstrap CI [0.26; 0.29], p < 0.001). A robust regression showed that for the BirdLife ranges, the overall accuracy had a significant effect on range overlap (*coef* = 0.54, *SE* = 0.015, *t* = 35.01, *p* <0.001). This means that decreasing accuracy in the BirdLife ranges decreases the spatial overlap with the IDW ranges and the expected estimates of AOH (Fig. 5)

**Figure 5.**
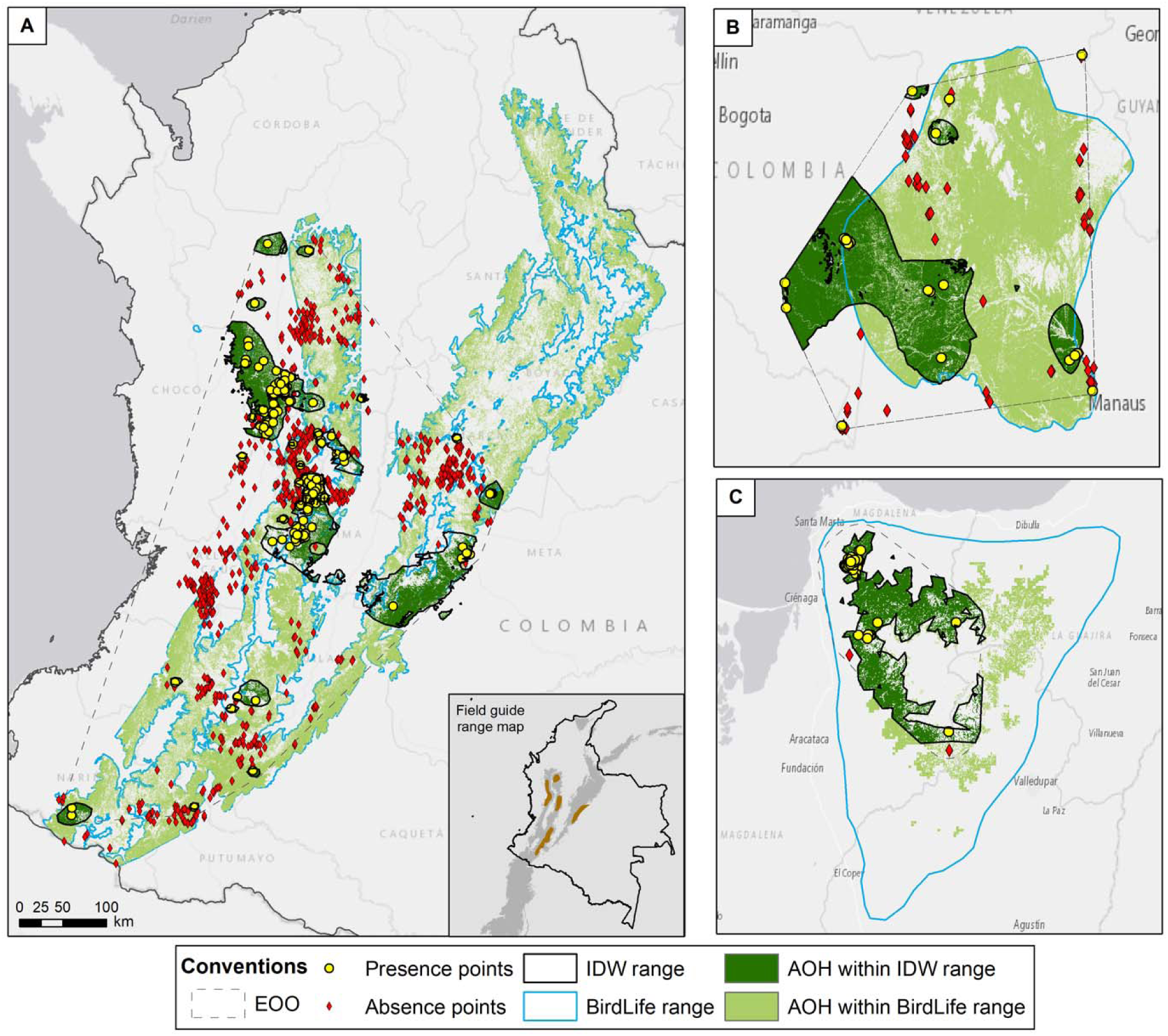
**A.** The Colombian endemic Yellow-eared parrot (*Ognorhynchus icterotis*) showed particularly poor spatial overlap between BirdLife ranges and IDW ranges, and respective estimated Area of Habitat (AOH). Note that BirdLife range extends the distribution north of the Eastern Andes where there are no records. On the inset, the expert-derived range map from Ayerbe-quiñones (2019) represented the species distribution much closer to the IDW range. **B.** For the Tawny-tufted Toucanet (*Selenidera nattereri*) in the Amazonian lowlands, the AOH derived from BirdLife is nearly three times larger than the estimated AOH derived from the IDW range. **C.** The BirdLife range for the Brown-rumped Tapaculo (*Scytalopus latebricola*) is a polygon covering the whole area of the Santa Marta mountains. In contrast, the IDW range restricted the AOH to the western side where observations have been recorded.

### 3. IUCN extinction risk assessments

For the 2,273 bird species analyzed, we found that based on criterion B the 2019 IUCN Red List (hereafter Red List) had 131 species in threatened categories whereas our preliminary assessment had 199, an increase of 51.9%. Moreover, there is only a moderate agreement between both criterion B assessments in terms of species assigned categories (Kendall’s *W* = 0.65, *p* < 0.001). We also identified 103 species as threatened that are currently classified as non-threatened categories (LC or NT) according to criteria B in IUCN (2019).

We analyzed the estimates of AOO and EOO derived from the occurrence data and those from the Red List. There was not a statistically significant difference in EOO values between both datasets (K-S test = 0.03, *p* = 0.122). Nevertheless, we identified 142 species that could be placed in threatened categories based on EOO thresholds, 66 of which are currently classified in non-threatened categories. Critically, only 36 species (1.6%) had a value of AOO listed in the Red List (all species had EOO values), whereas we identified 57 species that triggered AOO thresholds, 37 of which are currently considered as non-threatened.

We found 148 species that do not have any recorded information on their elevational limits (upper or lower) based on the data available on the Red List. Additionally, 886 species do not have a recorded lower elevation but have an upper limit, whereas 11 species do not have an upper limit but have a lower limit recorded. In total, 1045 (46%) have missing information regarding elevational limits. For the remaining 1,228 species that have complete elevational limits, 892 species (73%) have a narrower elevational range than the one found in our geospatial workflow, whereas 336 (27%) have a broader one. The difference in elevational range for both datasets was significant (K-S test = 0.30, *p* < 0.001).

## Discussion

We have developed an automated and reproducible geospatial workflow to map species distributions and assess extinction risk starting from primary biodiversity data, in a way that is consistent with the guidelines and criteria of the IUCN Red List. Using a dataset of 2,273 bird species in the Americas, 55% of all terrestrial birds found in the region, we have shown that our approach produced ranges that have reduced errors of omission, fewer errors of commission, and a higher accuracy when compared to expert-drawn range maps. Thus, it also provides a more reliable, transparent, and consistent way to derive estimates of Area of Habitat (AOH), the newly recommended measure for the IUCN Red List (Brooks et al. 2019).

Our workflow emphasized confirmed records and excluded potential distributional areas outside of each species EOO (Extent of Occurrence). Therefore, the generated ranges are especially useful for systematic conservation planning (Margules & Pressey 2000) where resources are directed to areas in which a species is known to be present. Unlike most expert-drawn range maps, we also excluded areas of known absence. Although absences always carry a high degree of uncertainty (Lobo et al. 2010) they are crucial for increasing the predictive accuracy of species ranges and therefore deriving better estimates of AOH (Elith et al. 2006; Mainali et al. 2020). The maps of AOH derived from our ranges could be valuable for conservationists in activities such as establishing protected areas or designing habitat corridors. On the other hand, given the low accuracy that we found on the expert-drawn range maps, we concur with Peterson et al. (2018) and do not recommend procedures that refine such maps to obtain estimates of AOH for guiding conservation efforts.

We have provided an objective and transparent way to produce species ranges, but we do not intend it for every scenario — there are no ‘silver bullets’ to represent species distributions for every purpose (Guillera-Arroita et al. 2015; Qiao et al. 2015). We do not recommend using our ranges for macroecological studies (*e.g*. species richness or endemism patterns) given that our predictions stay close to the presence points available and are limited by sampling bias. These biases might have influenced our findings that the BirdLife ranges were consistently larger than our generated ranges, but our findings are in line with previous research demonstrating substantial overestimation of range size from expert-drawn maps (Hurlbert & Jetz 2008; Jetz et al. 2008). Moreover, we also showed that range overestimation was driven by commission errors (false presences inside the range) that are prevalent in the BirdLife range maps. In contrast, our ranges markedly reduced commission errors and have a higher overall accuracy. Hence, our ranges are a reliable alternative for representing current knowledge on species distributions for conservation applications, with the advantage of being produced with an explicit and reproducible method.

Our study also highlighted the importance of incorporating geospatial data within the IUCN red listing process to improve species extinction risk estimates. We show with an automated assessment that when using occurrence data (GBIF), the number of potentially threatened species increased by 52% and identified up to 103 species likely threatened that are currently considered in non-threatened categories (LC or NT) based on criterion B of the 2019 IUCN Red List. We stress that this analysis is preliminary and is not intended to replace a full Red List assessment. Nevertheless, it does suggest that the IUCN Red List might be underestimating species extinction risk given the missing spatial information in key assessment parameters (EOO and AOO). For instance, while we derived AOO estimates for all the species in our dataset, less than 2% of species had a reported value of AOO in the IUCN Red List and 46% had missing elevational limits. Although the estimates of EOO between our methods and the Red List were not statistically different, it was sufficient to produce differences in the threat categorization of many species, with only moderate agreement between our preliminary conservation assessment and the 2019 IUCN Red List.

We recommend that the IUCN Red List incorporates more transparent and reproducible protocols, such as the one we have presented in this paper, to aid in mapping species distributions and increase transparency in species risk assessments. Our workflow could be useful to narrow down species that meet threatened criteria for further consultation with experts. Additional suggestions can be found in the literature - these include adopting species range model metadata standards (Merow et al. 2019), checklists to reproduce ecological niche models (Feng et al. 2019) and standards for biodiversity assessments (Araújo et al. 2019). These could help update the IUCN Rules of Procedure (IUCN 2016b) and Mapping Standards and Data Quality (IUCN Red List Technical Working Group 2019). Following suit, we suggest that all the spatial data used for assessments under the IUCN Red List should be made publicly available so that it can be fully understood, reviewed, and updated (Durant et al. 2017).

Our study demonstrated the utility and importance of a data-driven, reproducible approach to integrate mapping species distributions and extinction risk assessments in a framework compatible with the IUCN Red List. We have shown how our proposed workflow (1) assisted the red listing process by incorporating geospatial data and analysis to detect more potentially threatened species; (2) improved the accuracy of species ranges over expert-drawn range maps, and; (3) generated more reliable estimates of Area of Habitat (AOH) for local-scale planning and decision-making. In the urgency to evaluate the conservation status of many more species and implement measures to halt biodiversity loss (IUCN Species Survival Commission 2019), our workflow is a valuable addition to the toolkit of conservation practitioners and decision-makers because it is consistent, easily understood and widely applicable. Notwithstanding, we acknowledge the scarcity of spatial data is limiting in many groups and there are no easy shortcuts to address this shortfall (Rodrigues 2011). Only with more field explorations and carefully designed biological surveys, we will overcome the scarce knowledge on species occurrences and distributions around the globe for effective on-the-ground conservation actions (Wilson 2017).

## Acknowledgements

We thank S.L. Pimm and R. Huang for suggestions during the initial development of the geospatial workflow. We thank N. Hazzi and M. Di Marco for useful comments and edits. The first author thanks J. Meyer for encouragement to write the manuscript. This material is based upon work supported by the National Science Foundation under Award Numbers DBI-0735191, DBI-1265383, and DBI-1743442. URL: www.cyverse.org.

## Author contributions

R.D.P. conceived and developed the geospatial workflow, performed the analysis and drafted the manuscript. P.J.N., J.V-T, and A.P.J. provided critical input to the analysis and contributed with extensive edits. All authors gave final approval for publication.

## Data Availability

The bird species distribution maps of the world are available upon request to BirdLife International (http://datazone.birdlife.org/species/requestdis). Access to the IUCN Red List API is granted upon request (https://apiv3.iucnredlist.org/api/v3/token) and information is freely accessible at www.iucnredlist.org. Species occurrences can be gathered from its data providers through GBIF.org. The eBird observations and associated metadata can be obtained upon completion of a data request form (https://ebird.org/data/download). The SRTM v4.1 DEM is freely available for download from the CGIAR-CSI Consortium for Spatial Information (http://srtm.csi.cgiar.org/srtmdata/) The TanDEM-X Forest/Non-Forest Map - Global can be freely downloaded from (https://download.geoservice.dlr.de/FNF50/)

## Code Availability

The R code for the geospatial workflow will be made available upon reasonable request, previous to its deposition in an open-access repository with the peer-reviewed version of this study. Requests should be sent to the corresponding author.

